# Chemogenetic activation of astrocytes modulates sleep/wakefulness states in a brain region-dependent manner

**DOI:** 10.1101/2024.06.03.597103

**Authors:** Yuta Kurogi, Tomomi Sanagi, Daisuke Ono, Tomomi Tsunematsu

**Author notes:** **Address for correspondence:** Tomomi Tsunematsu, Ph.D. Department of Biology Faculty of Science Hokkaido University Sapporo 060-0810, Japan.

## Abstract

**Study objectives:** Astrocytes change their intracellular calcium (Ca^2+^) concentration during sleep/wakefulness states in mice. Furthermore, the Ca^2+^ dynamics in astrocytes vary depending on the brain region. However, whether alterations in intracellular Ca^2+^ concentration in astrocytes can affect sleep/wakefulness states and cortical oscillations in a brain region-dependent manner remain unclear.

**Methods:** The Ca^2+^ concentration in astrocytes was artificially increased using chemogenetics in mice. Astrocytes in the hippocampus and pons, which are 2 brain regions previously classified into different clusters based on their Ca^2+^ dynamics during sleep/wakefulness, were focused on to compare whether there are differences in the effects of astrocytes from different brain regions.

**Results:** The activation of astrocytes in the hippocampus significantly decreased the total time of wakefulness and increased the total time of sleep. This had minimal effects on cortical oscillations in all sleep/wakefulness states. On the other hand, the activation of astrocytes in the pons substantially suppressed rapid eye movement (REM) sleep in association with a decreased number of REM episodes, indicating strong inhibition of REM onset. Regarding cortical oscillations, the delta wave component during non-REM sleep was significantly enhanced.

**Conclusions:** These results suggest that astrocytes modulate sleep/wakefulness states and cortical oscillations. Furthermore, the role of astrocytes in sleep/wakefulness states appears to vary among brain regions.

**Statement of Significance:** Sleep is an instinctive behavior for many organisms. Recently, it has been reported that not only neurons, but also astrocytes, a type of glial cell, contribute to sleep/wakefulness states. Intracellular Ca^2+^ concentration, an indicator of astrocyte activity, fluctuates during sleep/wakefulness states. However, it is still unclear whether changes in Ca^2+^ concentration in astrocytes can affect sleep/wakefulness states. In this study, we utilized chemogenetics to activate astrocytes in mice. Our results showed that activation of astrocytes in the hippocampus causes decreased wakefulness, and that in the pons causes decreased REM sleep. Therefore, our findings demonstrate that the activation of astrocytes modulates sleep/wakefulness states in a brain region-dependent manner.

**Graphical Abstract:** 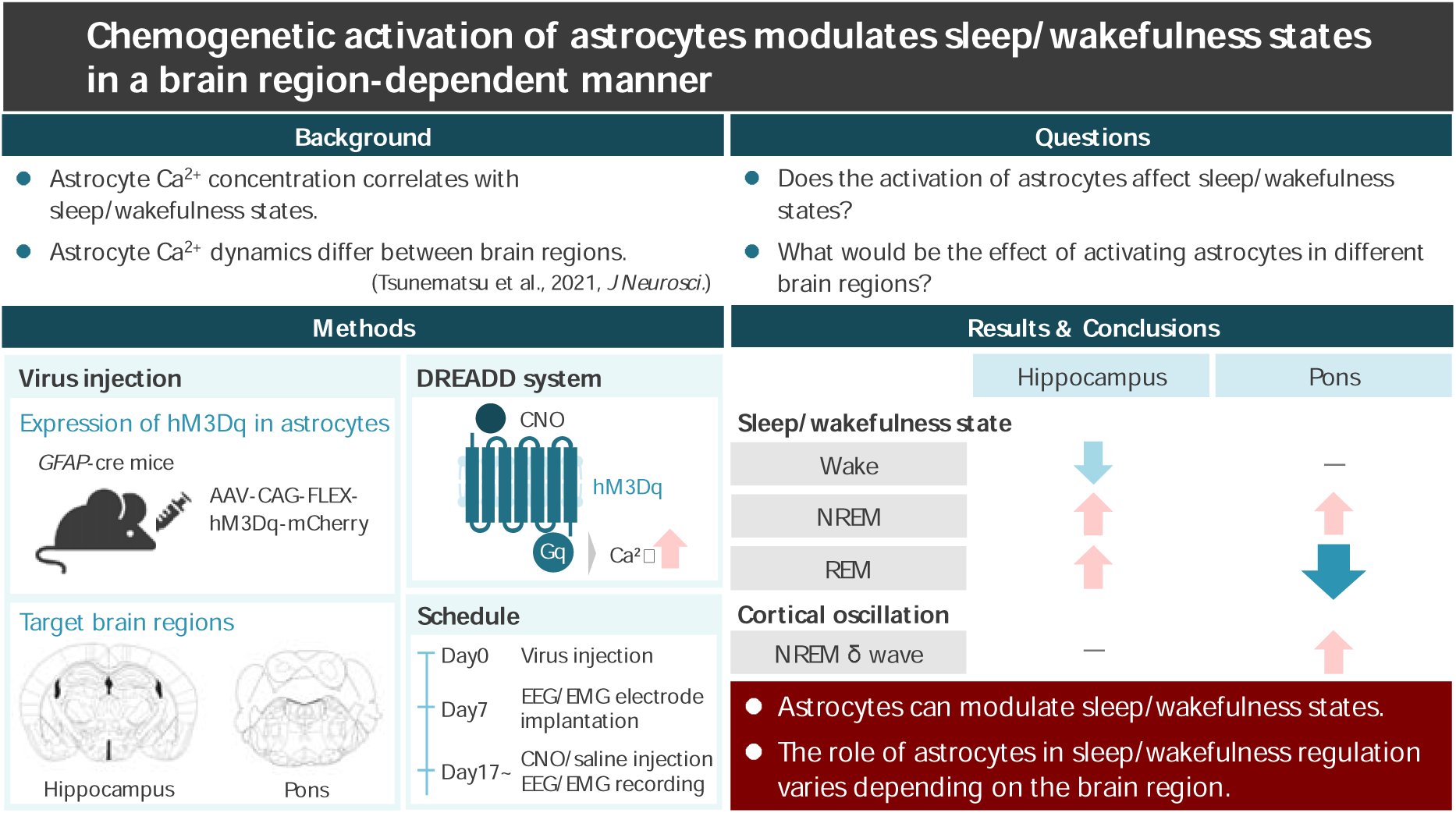

## Introduction

Astrocytes are one of the most abundant cell types in the brain. It is well known that astrocytes have housekeeping roles in brain function, contributing to ion buffering, neurotransmitter recycling, and metabolism regulation^1,2^. As astrocytes are non-excitable, all of their physiological functions are strongly affected by the dynamics of their intracellular Ca^2+^ concentration. Recently, a growing body of evidence has indicated that astrocytes play a crucial role in the regulation and physiological functions of sleep^3^. Astrocytes release a variety of sleep-promoting chemicals, such as cytokines, neurotrophins, prostaglandins, and purines^4-8^. In addition, several studies, including ours, have reported that intracellular Ca^2+^ concentrations in astrocytes substantially fluctuate through sleep/wakefulness states^9-11^. The intracellular Ca^2+^ concentration of astrocytes is highest during wakefulness and lowest during sleep, although this degree varies depending on the brain region^11^. These results indicate that astrocytes in different brain regions have different functions in sleep/wakefulness states.

Previous studies have suggested that intracellular Ca^2+^ dynamics in astrocytes play a crucial role, particularly in the regulation of rapid eye movement (REM) sleep. A decrease in intracellular Ca^2+^ concentrations in astrocytes occurs during sleep, and this decrease is more pronounced during REM sleep than during non-REM (NREM) sleep^9-11^. The reduction of inositol 1,4,5-trisphosphate in astrocytes throughout the brain results in a constant suppression of Ca^2+^ release from intracellular Ca^2+^ stores, and alters the duration and strength of theta waves during REM sleep^12^.

In the present study, we focused on 2 brain regions that play important roles in REM sleep. The pons, which contains various neurons that are involved in the regulation of REM sleep^13-15^, and the hippocampus, which generates theta waves, the electroencephalogram (EEG) characteristic of REM sleep^16,17^. Moreover, in our previous study, we showed that astrocytes in the hippocampus and the pons are classified into various different clusters based on their intracellular Ca^2+^ dynamics during sleep/wakefulness states^11^. We therefore suspected that they might have different roles in sleep/wakefulness modulation. Therefore, in this study, we investigated whether artificially increasing the intracellular Ca^2+^ concentration in astrocytes affects sleep/wakefulness states and cortical oscillations in mice. To answer these questions, we utilized a chemogenetics approach. Our results showed that increasing the intracellular Ca^2+^ concentration in hippocampal astrocytes decreased wakefulness and increased sleep. On the other hand, increasing the intracellular Ca^2+^ concentration in pons astrocytes substantially suppressed REM sleep. In addition, the delta wave component of the EEG during NREM sleep was enhanced. These results suggest that astrocyte activity regulates sleep/wakefulness states, and that different brain regions play different roles in sleep/wakefulness regulation.

## Methods

### Mice

All experimental procedures involving mice were approved by the Animal Care and Use Committee of Tohoku University (study approval no.: 2019LsA-018) and Hokkaido University (study approval no.: 23-0117), and were conducted in accordance with the National Institute of Health guidelines. All efforts were made to minimize animal suffering and discomfort, and to reduce the number of animals used. Mice were housed under a controlled 12 h/12 h light/dark cycle (light on hours: 8:00–20:00). Mice had *ad libitum* access to food and water. Glial fibrillary acidic protein (*GFAP*)-cre mice on a C57BL/6J background (024098, The Jackson Laboratory) were used to chemogenetically increase the intracellular calcium concentration in their astrocytes. The following polymerase chain reaction (PCR) primer sets were used for mouse genotyping: GFAPcre Fw (5′-TCCATAAAGGCCCTGACATC-3′) and GFAPcre Rv (5′-TGCGAACCTCATCACTCGT-3′). A total of 16 male *GFAP*-cre mice were used. For chemogenetics recordings, 12 mice were used. For cFos immunohistochemistry studies, 4 mice were used.

### Viruses

Adeno-associated viruses (AAVs) were generated using a triple-transfection, helper-free method, and purified following previously established procedures^18^. In brief, human embryonic kidney 293A cells (Invitrogen) were transfected with pHelper, pAAV5, and pAAV-CAG-FLEX-hM3Dq-mCherry using the standard calcium phosphate method. Three days after transfection, the cells were collected and suspended in phosphate buffer saline (PBS) containing 1 mM MgCl_2_. After 2 freeze-thaw cycles, the cell lysate was treated with benzonase nuclease (Merck) at 37 °C for 30 minutes, followed by centrifugation twice at 15,000 *g* for 10 minutes each. The supernatant containing the viruses was collected for further use. The titer of the AAV vector was determined by quantitative PCR: *AAV_5_-CAG-FLEX-hM3Dq-mCherry*, 2 × 10^12^. The final purified viruses were stored at −80 °C.

### Surgical procedures

Male *GFAP*-cre mice of approximately 3 to 6 months old were used (111.7 ± 6.3 days of age). Stereotaxic surgery was performed under anesthesia with pentobarbital (5 mg/kg, intraperitoneal [i.p.] injection as induction) and with isoflurane (1%–2% for maintenance) using a vaporizer for small animals (Bio Research Center), with the mice positioned in a stereotaxic frame (Narishige). For chemogenetics recordings, *AAV_5_-CAG-FLEX-hM3Dq-mCherry* was injected into the hippocampus (1.7 mm posterior, ± 0.8 mm lateral, and a depth of 2.0 mm from bregma), or pons (5.3 mm posterior, ± 1.0 mm lateral, and a depth of 4.0 mm from bregma) of *GFAP*-cre mice in a volume of 200 nL and at a flow rate of 20 nL per min using a Hamilton 10 μL syringe. The needle was kept in place for 10 min after the injection. The injection volume and flow rate were controlled by a stereotaxic injector (Legato 130, Muromachi Kikai). One week after AAV injection, surgery for EEG/electromyogram (EMG) recordings were performed. Three bone screws were implanted on the skull as electrodes for cortical EEGs, and twisted wires (AS633, Cooner wire) were inserted into the neck muscle as an electrode for EMGs. Another bone screw was implanted into the cerebellum as a ground. All electrodes were connected to a pin socket and fixed to the skull with dental cement. After the surgery, the mice were left to recover for at least 7 days, and then transferred to sleep recording chambers.

### Chemogenetics

Clozapine N-oxide (CNO) (ab141704, Abcam) was dissolved in saline to 1 mg/mL. CNO or saline was administered by i.p. injection to each mouse (0.3 mL/30 g body weight) at Zeitgeber time (ZT) 1. Each mouse received 3 saline and 3 CNO administrations in a random manner, at intervals of 3 days.

### *In vivo* sleep/wakefulness recording using freely moving mice

Continuous EEG and EMG recordings were conducted through a slip ring (SPM-35-8P-03, HIKARI DENSHI), which was designed so that the movement of the mice was unrestricted. EEG and EMG signals were amplified (AB-610J, Nihon Koden), filtered (EEG, 0.75–20 Hz; EMG, 20–50 Hz), digitized at a sampling rate of 128 Hz, and recorded using SleepSign software version 3 (Kissei Comtec). Mouse behavior was monitored through a charge coupled device video camera and recorded on a computer synchronized with EEG and EMG recordings using the SleepSign video option system (Kissei Comtec).

### Immunohistochemistry

To confirm the specific expression of hM3Dq in astrocytes, mice were deeply anesthetized with isoflurane, and perfused sequentially with 20 mL of chilled PBS and 20 mL of chilled 4% paraformaldehyde in PBS (Nacalai Tesque). To confirm the activation of astrocytes via CNO, mice were deeply anesthetized 1.5 hours after CNO or saline injection, and perfused in a similar way. The brains were removed and immersed in the above fixation solution overnight at 4 °C, and then immersed in 30% sucrose in PBS for at least 2 days. The brains were quickly frozen in embedding solution (Sakura Finetek), and cut into coronal sections using a cryostat (CM1850 or CM3050S, Leica) at a thickness of 40 µm. For astrocyte immunostaining, brain sections were incubated with an anti-GFAP antibody (1:2,000; G3893, Sigma) overnight at 4 °C. Then, the sections were incubated with CF488A donkey anti-mouse immunoglobulin G (IgG) (1:1,000; 20014-1, Nacalai Tesque) for 2 hours at room temperature. For microglia immunostaining, sections were incubated with an anti-ionized calcium-binding adapter protein 1 (Iba1) antibody (1:1,000; 019-19741, FUJIFILM Wako Pure Chemical) overnight at 4 °C. Then, the sections were incubated with CF488A donkey anti-rabbit IgG (1:1,000; 20015-1, Nacalai Tesque) for 2 hours at room temperature. To visualize neurons, Nissl staining was performed. The brain sections were incubated with NeuroTrace Green Fluorescent Nissl Stain (1:300; N21480, Invitrogen) for 1 hour at room temperature. For cFos immunostaining, brain sections were incubated with an anti-cFos antibody (1:5,000; 226008, Synaptic Systems GmbH) overnight at 4 °C. Then, the sections were incubated with CF488A donkey anti-rabbit IgG (1:1,000; 20015-1, Nacalai Tesque) for 2 hours at room temperature. All sections were incubated with 4’,6-diamidino-2-phenylindole (DAPI) (1:1,000; D523, Dojindo) for 30 min at room temperature to visualize nuclei. The sections were mounted onto APS-coated slides, coverslipped with 50% glycerol in PBS, and observed using a fluorescence microscope (BZ-X800L or BZ-9000, Keyence).

### Sleep scoring and EEG analysis

Polysomnographic recordings were automatically scored offline, with each epoch scored as wakefulness, NREM sleep, or REM sleep by SleepSign software (Kissei Comtec) in 4-sec epochs, in accordance with standard criteria^19,20^. All vigilance state classifications assigned by SleepSign were confirmed visually. The same individual, blinded to the experimental condition, scored all EEG/EMG recordings. Spectral analysis of the EEGs was performed by Fast Fourier transform, which yielded a power spectral profile with a 1-Hz resolution divided into delta (1−5 Hz), theta (6−10 Hz), alpha (10−13 Hz), and beta (13−25 Hz) waves.

### Statistical analysis

Data are presented as the mean ± standard error of the mean (SEM) unless otherwise stated. Statistical analyses were performed using MATLAB software (MathWorks). Data were analyzed by the paired *t*-test. A *p*-value of less than 0.05 was considered to indicate a statistically significant difference between groups.

## Results

### Specific expression of hM3Dq in hippocampal astrocytes

To study the role of hippocampal astrocytes in sleep/wakefulness regulation, we introduced a designer receptor exclusively activated by designer drug (DREADD) in astrocytes. To increase intracellular Ca^2+^ concentration in astrocytes, we bilaterally injected the excitatory chemogenetic virus *AAV-CAG-FLEX-hM3Dq-mCherry* into the hippocampus of *GFAP*-cre mice (Figure 1A). The specific expression of hM3Dq in astrocytes was confirmed by immunohistochemical staining using the astrocyte-specific marker GFAP. The merged images show that hM3Dq-mCherry was exclusively observed in hippocampal astrocytes (Figure 1B). Iba1 staining for microglia, and NeuN staining for neurons showed no staining of the hM3Dq-mCherry-positive cells (Figure 1C and 1D), indicating no ectopic expression.

**Figure 1.**
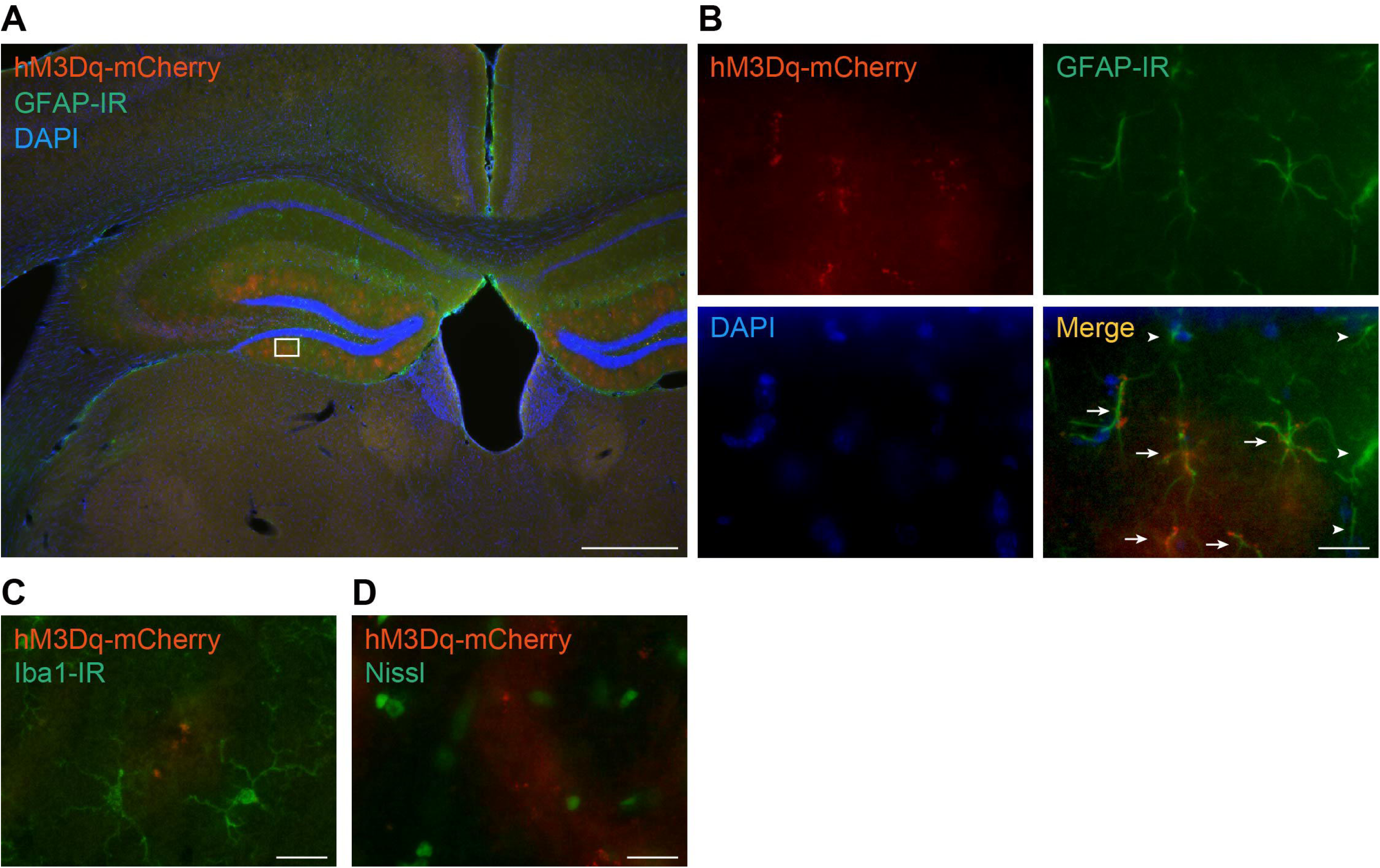
Specific expression of hM3Dq in hippocampal astrocytes. (A) Expression pattern of hM3Dq in the hippocampus of *GFAP*-cre mice. Red, hM3Dq-mCherry; green, GFAP-IR astrocytes; blue, DAPI. Scale bar: 500 μm. (B) Higher magnifications of the square region in (A). Arrows indicate GFAP-IR astrocytes expressing hM3Dq. Arrowheads indicate GFAP-IR astrocytes not expressing hM3Dq. Scale bar: 10 μm. (C) Fluorescence immunostaining using an anti-Iba1 antibody. Red, hM3Dq-mCherry; green, Iba1-IR microglia. Scale bar: 10 μm. (D) Nissl staining. Red, hM3Dq-mCherry; green, Nissl-positive neurons. Scale bar: 10 μm

### Chemogenetic activation of hippocampal astrocytes decreases wakefulness and increases sleep

First, to confirm whether the i.p. administration of CNO (1 mg/kg) induces the activation of astrocytes, we performed immunostaining of cFos, which is an immediately early gene, as a marker of cellular activation. The mice were injected with CNO or saline, and their brains were collected after 90 min. Immunohistochemical analyses demonstrated that the mice with chemogenetic activation of astrocytes had increased cFos expression in the hippocampus compared with saline-treated control mice (Figure 2A).

**Figure 2.**
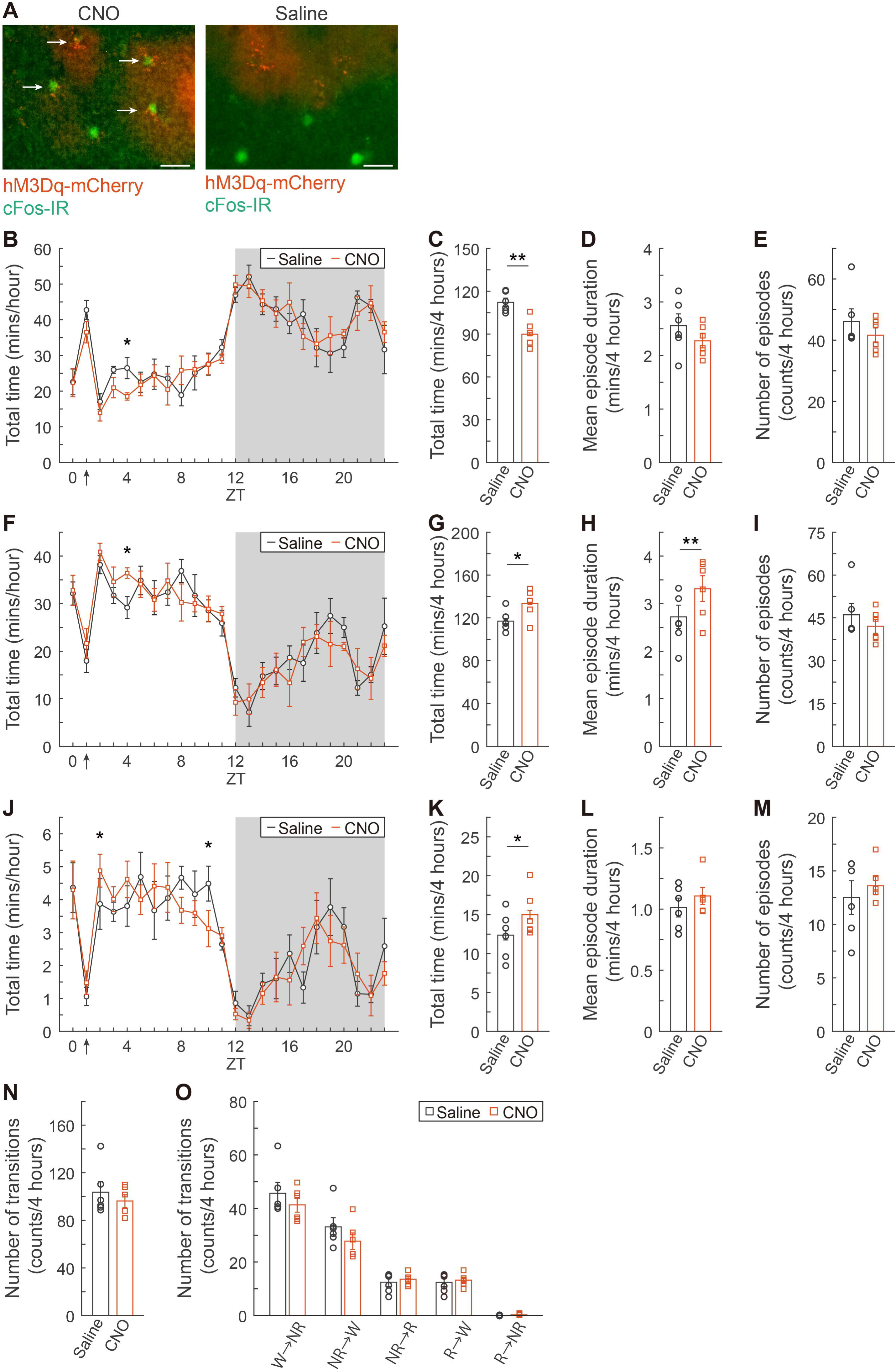
Effects of the chemogenetic activation of hippocampal astrocytes on sleep/wakefulness. (A) CNO administration to mice expressing hM3Dq in their hippocampal astrocytes resulted in increased cFos expression (left) in the astrocytes (arrows) compared with saline-treated mice (right). Red, hM3Dq-mCherry; green, cFos-IR astrocytes. Scale bar: 10 μm. (B, F, and J) Hourly amounts of wakefulness (B), NREM sleep (F), and REM sleep (J) after i.p. injection of saline or CNO (1 mg/mL) at ZT1 (arrows). The gray background indicates the dark period. (C, G, and K) Total amount of wakefulness (C), NREM sleep (G), and REM sleep (K) in the 4 hours after i.p. injection of saline or CNO. (D, H, and L) Mean episode duration of wakefulness (D), NREM sleep (H), and REM sleep (L). (E, I, and M) Number of episodes of wakefulness (E), NREM sleep (I), and REM sleep (M). (N) The number of transitions between sleep/wakefulness states during the 4 hours after saline or CNO administration. (O) The number of transitions between sleep/wakefulness states during the 4 hours after saline or CNO administration. Values are represented as means ± SEM; *, *p* < 0.05. **, *p* < 0.01

To determine the sleep/wakefulness states of mice during chemogenetic activation of hippocampal astrocytes, mice were chronically implanted with EEG and EMG electrodes in freely moving conditions. *GFAP*-cre mice expressing hM3Dq specifically in hippocampal astrocytes were injected i.p. with CNO (1 mg/kg) or with saline as a control at ZT1 (n = 6), and their hourly amount of sleep/wakefulness was analyzed (Figure 2B, 2F, and 2J). The activation of hippocampal astrocytes significantly decreased the total time of wakefulness at ZT4 compared with control (*t(5)* = –3.31, *p* < 0.05, paired *t*-test; Figure 2B) in association with decreased total time of NREM sleep at the same time (*t(5)* = 3.39, *p* < 0.05, paired *t*-test; Figure 2F). The total time of REM sleep was significantly increased at ZT2 and decreased at ZT10 (ZT2: *t(5)* = 2.87, *p* < 0.05, paired *t*-test; ZT10: *t(5)* = –3.09, *p* < 0.05, paired *t*-test; Figure 2J). During the 4 hours post-injection period (from ZT1 to ZT4), the total time of wakefulness significantly decreased (*t(5)* = –5.03, *p* < 0.01, paired *t*-test; Figure 2C). In contrast, the total time of NREM sleep and REM sleep significantly increased (NREM sleep: *t(5)* = 2.93, *p* < 0.05, paired *t*-test; REM sleep: *t(5)* = 2.61, *p* < 0.05, paired *t*-test; Figure 2G and 2K). There was a significant increase in the mean episode duration of NREM sleep (*t(5)* = 4.24, *p* < 0.01, paired *t*-test; Figure 2H), but not for wakefulness or REM sleep (Figure 2D and 2L). There were no significant changes in the number of episodes across all sleep/wakefulness states (Figure 2E, 2I, and 2M). Transition frequencies during all sleep/wakefulness states and in each state did not change in the 4-hour post-injection period (Figure 2N, and 2O). These results demonstrated that the activation of astrocytes in the hippocampus induces a decrease in wakefulness and an increase in sleep.

### Hippocampal astrocyte activation minimally affects cortical oscillations

As the activation of hippocampal astrocytes caused changes in sleep/wakefulness states, we next analyzed cortical oscillations based on the EEG in each sleep/wakefulness state for 4 hours after CNO injection (n = 6). EEG power values in the 1-Hz bin were normalized for a total of 1 to 20 Hz EEG power during each of wakefulness, NREM sleep, and REM sleep (Figure 3A, 3C, and 3E, respectively). The activation of hippocampal astrocytes significantly increased the normalized power at 4 Hz during NREM sleep compared with control (*t(5)* = 2.88, *p* < 0.05, paired *t*-test; Figure 3C), but there was no change during wakefulness and NREM sleep (Figure 3A, and 3E). We further analyzed the EEG in frequency bands, with 1 to 5 Hz as delta waves, 6 to 10 Hz as theta waves, 10 to 13 Hz as alpha waves, and 13 to 20 Hz as beta waves (Figure 3B, 3D, and 3F, respectively). No significant differences, however, were found compared with control. These results indicate that the activation of hippocampal astrocytes has minimal effects on the EEG during sleep/wakefulness states.

**Figure 3.**
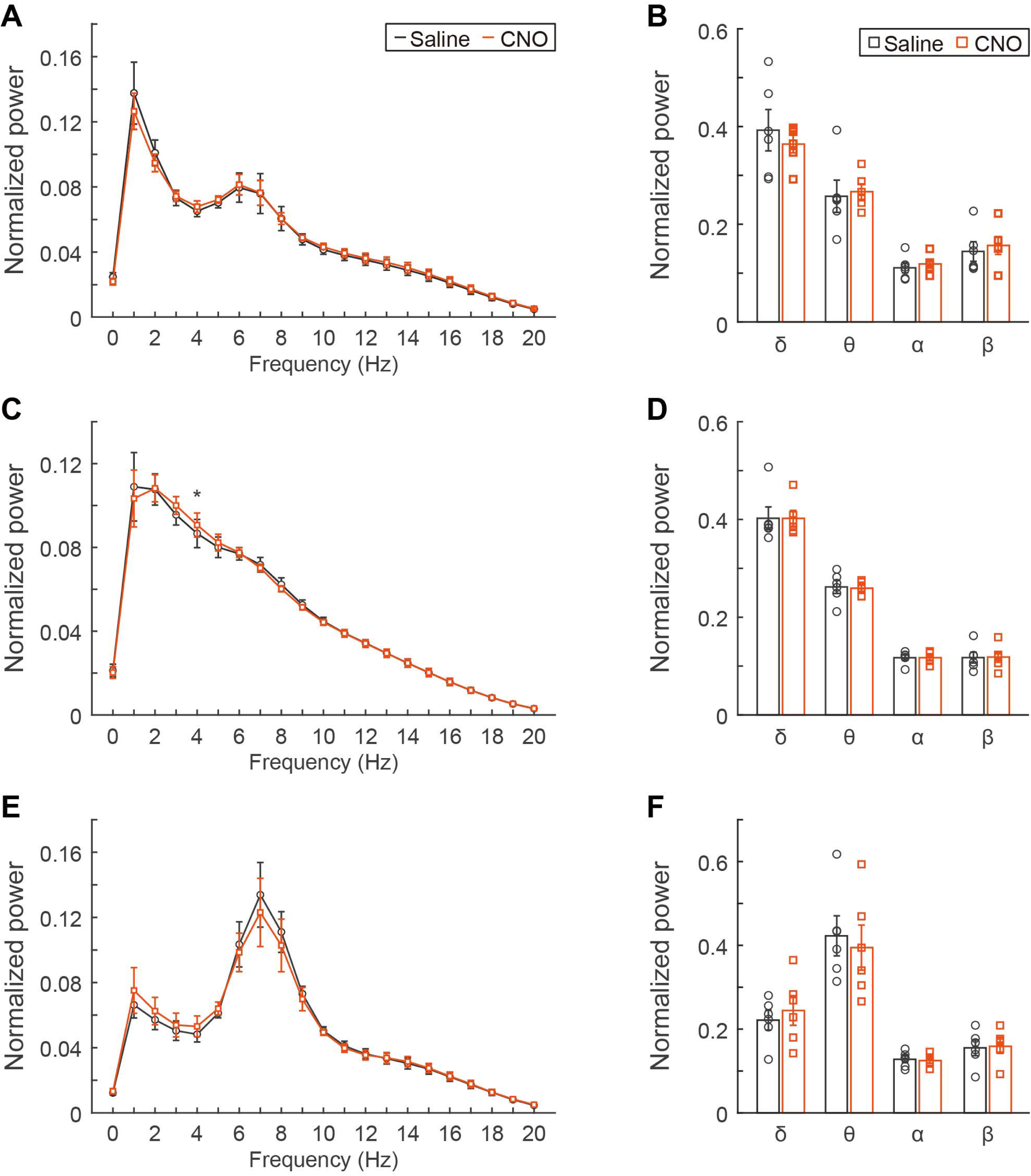
Effect of the chemogenetic activation of hippocampal astrocytes on EEG during sleep/wakefulness states. (A, C, and E) Average normalized EEG power spectra during wakefulness (A), NREM sleep (B), and REM sleep (C) for the 4 hours immediately after saline or CNO (1 mg/mL) administration at ZT1. (B, D, and F) Comparison of normalized delta (1–5 Hz) power, theta (6–10 Hz) power, alpha (10–13 Hz) power, and beta (13–20 Hz) power during wakefulness (B), NREM sleep (D), and REM sleep (F). Values are represented as means ± SEM; *, *p* < 0.05

### Specific expression of hM3Dq in pons astrocytes

Next, to study the role of pons astrocytes in sleep/wakefulness regulation, intracellular Ca^2+^ concentration was increased in astrocytes in the pons as well as in the hippocampus, using chemogenetics (Figure 4A). The specific expression of hM3Dq in astrocytes was confirmed by immunohistochemical staining of the astrocyte-specific marker GFAP. The merged images show that hM3Dq-mCherry was exclusively observed in pons astrocytes (Figure 4B). None of the microglia and neurons stained with Iba1 and NeuN, respectively, were hM3Dq-mCherry-positive (Figure 4C and 4D, respectively), indicating the absence of ectopic expression.

**Figure 4.**
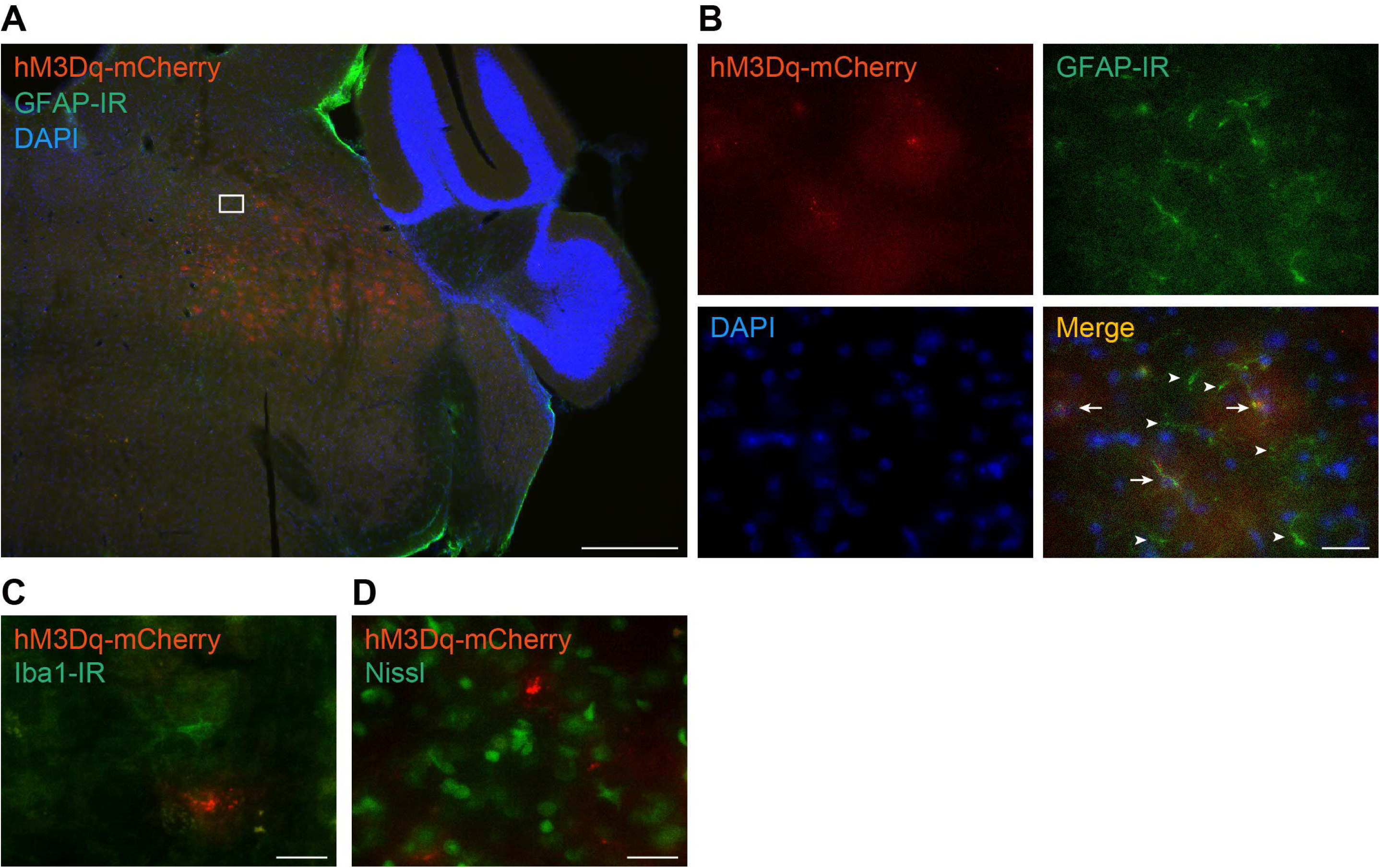
Specific expression of hM3Dq in astrocytes in the pons. (A) Expression pattern of hM3Dq in the pons in *GFAP*-cre mice. Red, hM3Dq-mCherry; green, GFAP-IR astrocytes; blue, DAPI. Scale bar: 500 μm. (B) Higher magnification images of the square region in (A). Arrows indicate GFAP-IR astrocytes expressing hM3Dq. Arrowheads indicate GFAP-IR astrocytes not expressing hM3Dq. Scale bar: 10 μm. (C) Fluorescence immunostaining with an anti-Iba1 antibody. Red, hM3Dq-mCherry; green, Iba1-IR microglia. Scale bar: 10 μm. (D) Nissl staining. Red, hM3Dq-mCherry; green, Nissl-positive neurons. Scale bar: 10 μm

### Chemogenetic activation of pons astrocytes substantially decreases REM sleep

cFos expression was analyzed immunohistochemically to determine whether the i.p. administration of CNO induces the activation of astrocytes in the pons as well as in the hippocampus. We confirmed that mice with chemogenetic activation of their astrocytes showed increases cFos expression in the pons compared with saline-treated control mice (Figure 5A).

**Figure 5.**
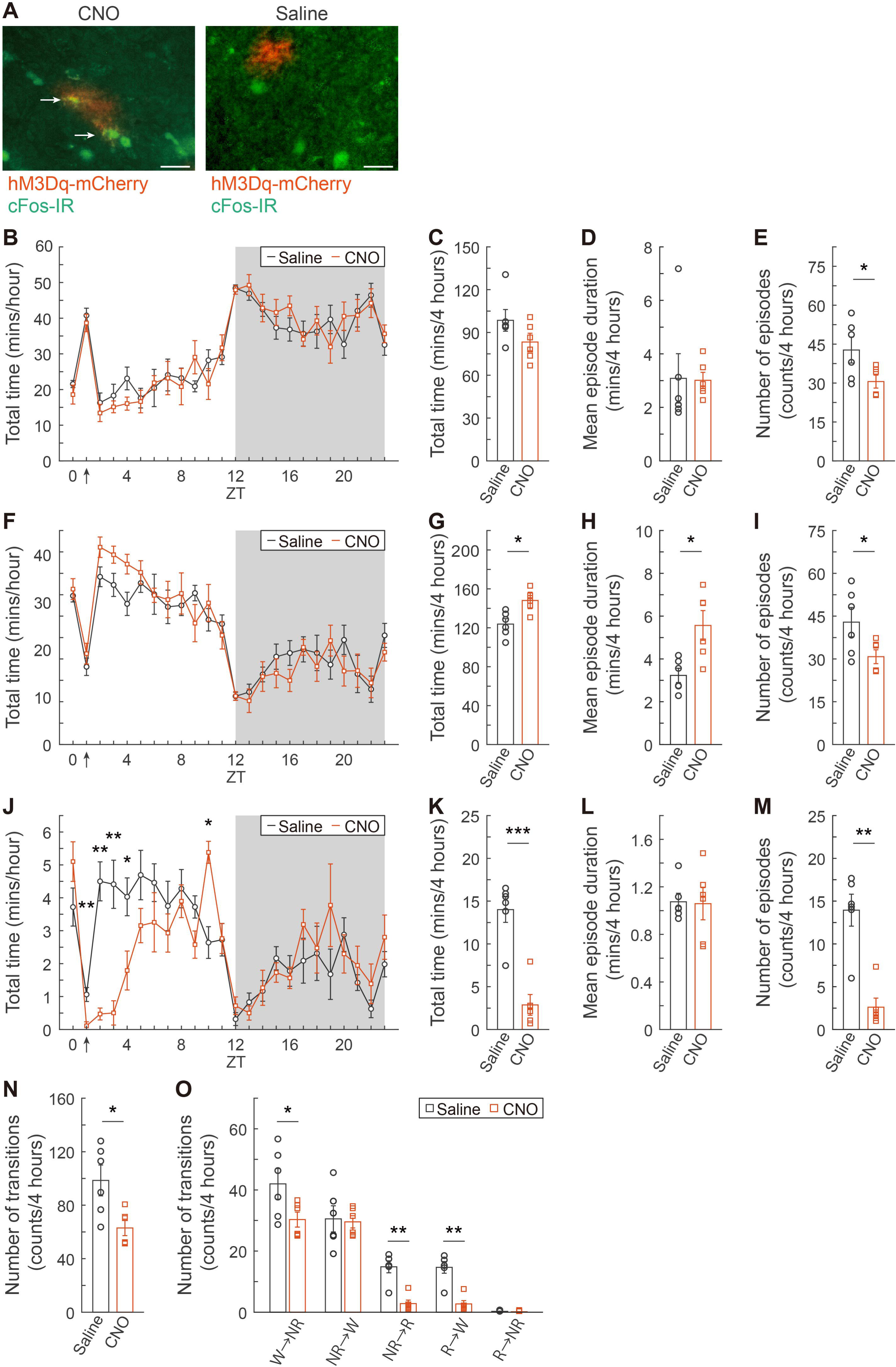
Effects of the chemogenetic activation of pons astrocytes on sleep/wakefulness. (A) CNO (1 mg/mL) administration to mice expressing hM3Dq in their pons astrocytes resulted in an increase in cFos expression (left) in these astrocytes (arrows) compared with saline-treated mice (right). Red, hM3Dq-mCherry; green, cFos-IR astrocytes. Scale bar: 10 μm. (B, F, and J) Hourly amounts of wakefulness (B), NREM sleep (F), and REM sleep (J) after i.p. injection of saline or CNO (1 mg/mL) at ZT1 (arrows). Gray background indicates the dark period. (C, G, and K) Total amounts of wakefulness (C), NREM sleep (G), and REM sleep (K) during the 4 hours after the i.p. injection of saline or CNO (1 mg/mL). (D, H, and L) Mean episode duration of wakefulness (D), NREM sleep (H), and REM sleep (L). (E, I, and M) Number of episodes of wakefulness (E), NREM sleep (I), and REM sleep (M). (N) The number of transitions between all states during the 4 hours after saline or CNO administration. (O) The number of transitions between each sleep/wakefulness state during the 4 hours after saline or CNO administration. Values represent means ± SEM; *, *p* < 0.05. **, *p* < 0.01, ***, *p* < 0.001

To determine the physiological role of activated astrocytes in the pons and whether they have a different role in sleep/wakefulness states compared with hippocampal astrocytes, *GFAP*-cre mice expressing hM3Dq specifically in pons astrocytes were injected i.p. with CNO (1 mg/kg) or with saline as a control at ZT1 (n = 6). Sleep/wakefulness states were recorded using freely moving mice. The hourly amounts of sleep/wakefulness states for 24 hours showed that REM sleep was strongly suppressed after CNO injection (Figure 5J), whereas there was no significant change in both wakefulness and NREM sleep (Figure 5B and 5F, respectively). The activation of pons astrocytes significantly decreased the total time of REM sleep during the 4 hours after CNO administration compared with control astrocytes (*t(5)* = –7.09, *p* < 0.001, paired *t*-test; Figure 5K). The total time of NREM sleep was significantly increased (*t(5)* = 2.88, *p* < 0.05, paired *t*-test; Figure 5G). On the other hand, there was no difference in the total time of wakefulness (Figure 5C). In terms of wakefulness episodes, there was no significant change in duration, but a significant reduction in number (*t(5)* = –2.74, *p* < 0.05, paired *t*-test; Figure 5D–E). During NREM sleep, mean episode duration significantly increased (*t(5)* = 4.80, *p* < 0.01, paired *t*-test), in association with a decrease in the number of episodes (*t(5)* = –2.59, *p* < 0.05, paired *t*-test; Figure 5H–I). In contrast, regarding REM sleep episodes, mean duration did not change, but the number of episodes significantly decreased (*t(5)* = –6.51, *p* < 0.01, paired *t*-test; Figure 5L–M). A significant decrease in transition frequency was observed across all sleep/wakefulness states (*t(5)* = –3.34, *p* < 0.05, paired *t*-test; Figure 5N). The transition frequency to each state was further analyzed in detail (Figure 5O). The activation of pons astrocytes via CNO injection induced a significant decrease in transition number from wakefulness to NREM sleep (*t(5)* = –2.58, *p* < 0.05, paired *t*-test), from NREM sleep to REM sleep (*t(5)* = –6.36, *p* < 0.01, paired *t*-test), and from REM sleep to wakefulness (*t(5)* = –6.36, *p* < 0.01, paired *t*-test). Taken together, these findings indicate that the activation of pons astrocytes reduces REM sleep by strongly inhibiting its onset.

### Activation of pons astrocytes increases delta waves during NREM sleep

Next, to analyze the effect of the activation of pons astrocytes on the cortical oscillations during sleep/wakefulness states, a spectral analysis of EEG was performed. EEG power was normalized as in the analysis of the activation of hippocampal astrocytes (Figure 3). Despite the results that the activation of pons astrocytes strongly suppressed REM sleep, unexpectedly, EEG power was substancially altered in NREM sleep, but not in wakefulness and REM sleep (Figure 6A, 6C, and 6E, respectively): the normalized power of EEG between 2 to 5 Hz during NREM sleep in these mice was significantly increased, and the normalized power of EEG between 7 to 20 Hz was significantly decreased compared with control mice (Figure 6C). With respect to each frequency band, during NREM sleep, delta waves were significantly higher in CNO-treated mice than those in the control mice, whereas theta, alpha, and beta waves were significantly lower (delta: *t(5)* = 3.83, *p* < 0.05, paired *t*-test; theta: *t(5)* = –3.43, *p* < 0.05, paired *t*-test; alpha: *t(5)* = –4.96, *p* < 0.01, paired *t*-test; beta: *t(5)* = –5.57, *p* < 0.01, paired *t*-test; Figure 6D). On the other hand, there were no significant differences in wakefulness and REM sleep (Figure 6B, and 6F). In conclusion, we demonstrated that the activation of astrocytes in the pons increases the low-frequency components in EEG during NREM sleep in mice.

**Figure 6.**
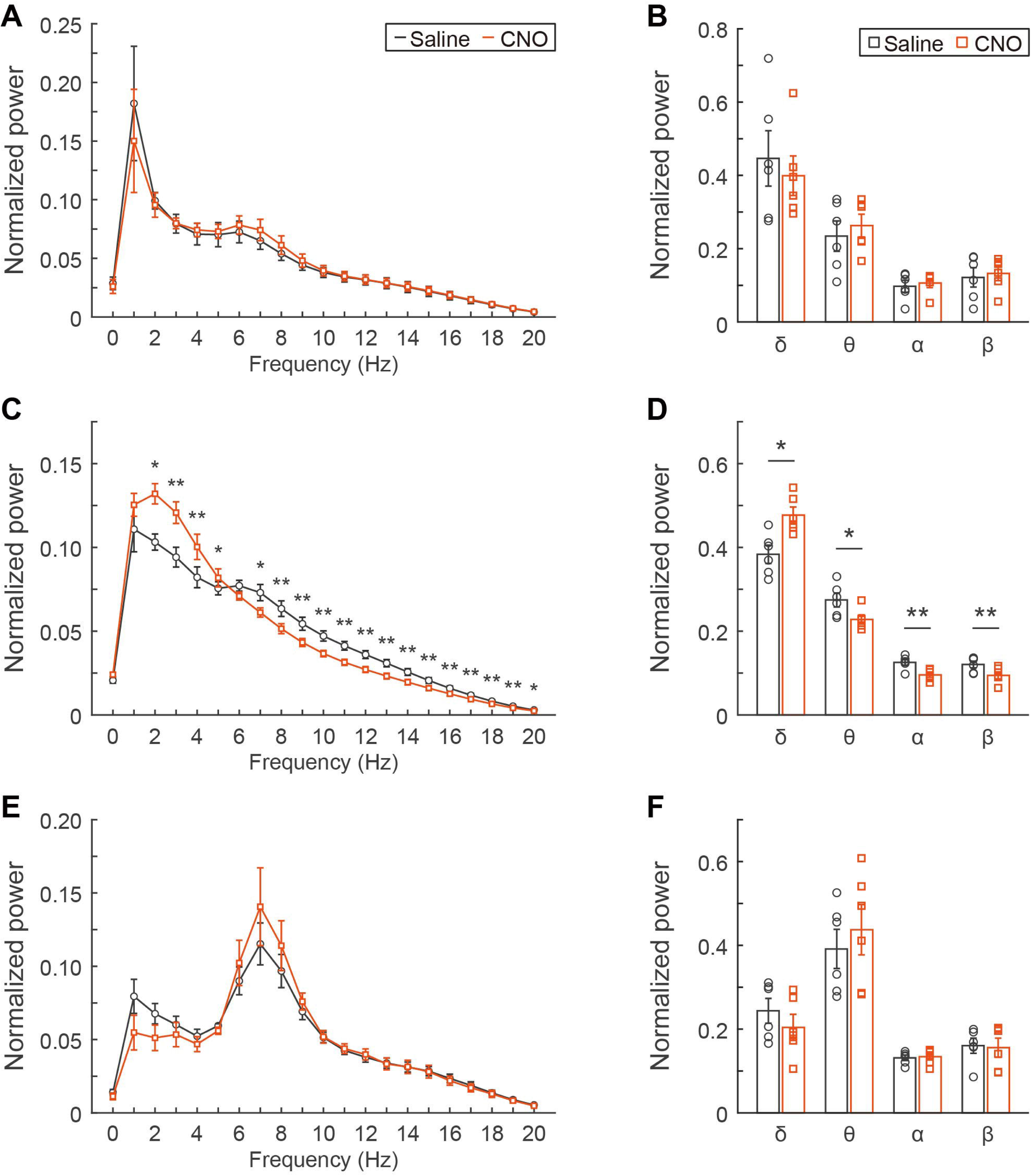
Effect of the chemogenetic activation of pons astrocytes on EEG during sleep/wakefulness states. (A, C, and E) Average normalized EEG power spectra during wakefulness (A), NREM sleep (B), and REM sleep (C) for 4 hours after saline or CNO (1 mg/mL) administration at ZT1. (B, D, and F) Comparison of normalized delta (1–5 Hz) power, theta (6–10 Hz) power, alpha (10–13 Hz) power, and beta (13–20 Hz) power during wakefulness (B), NREM sleep (D), and REM sleep (F). Values represent means ± SEM; *, *p* < 0.05.**, *p* < 0.01

## Discussion

In the present study, we analyzed the sleep/wakefulness states and cortical oscillations in mice under conditions of increased intracellular Ca^2+^ concentrations in hippocampal and pons astrocytes using chemogenetics. The activation of hippocampal astrocytes induced a significant decrease in the total time of wakefulness, and a significant increase in the total time of NREM sleep and REM sleep. There was almost no effect on cortical oscillations in any of the sleep/wakefulness states. On the other hand, activation of pons astrocytes substantially decreased the total time of REM sleep, accompanied with suppression of REM occurrence. There was a significant reduction in the total time of NREM sleep, but no effect on the total time of wakefulness. Moreover, the delta component during NREM sleep was significantly increased.

### Activation of astrocytes modulates sleep/wakefulness states

Previous studies, including ours, demonstrated that the dynamics of intracellular Ca^2+^ concentration in astrocytes is associated with sleep/wakefulness states^9-11^. The causal association between the activity of astrocytes and sleep/wakefulness states has not been completely clarified to date, although several studies using chemogenetics techniques have been reported in the previous few years^21-24^. The use of chemogenetics has considerable side effects on sleep studies; it has been reported that the administration of CNO to mice that do not express DREADD can alter the total duration of NREM and REM sleep, as well as the number of episodes and episode duration^25,26^. However, such side effects are observed only when CNO is administered at high doses (5 or 10 mg/kg).

At the same dose with which CNO was administered in the present study (1 mg/kg), any effects on sleep/wakefulness states were not observed^25^. In other words, it is clear that the changes in sleep/wakefulness states observed in our present study were a result of the activation of astrocytes by CNO administration, and not the CNO administration itself. Therefore, astrocytes are able to modulate sleep/wakefulness states.

We compared the effects of activation of hippocampal and pons astrocytes on sleep/wakefulness states, to clarify whether there were any differences. Our results demonstrated that the activation of hippocampal astrocytes induced a moderate decrease in wakefulness and an increase in sleep. This result is inconsistent with a report that optogenetic activation of hippocampal astrocytes by expressing channelrhodopsin-2 (ChR2) does not change sleep/wakefulness states^27^. This discrepancy may be owing to differences in the experimental design between chemogenetics and optogenetics. Astrocytes express G-protein-coupled receptors, but not cation channels, such as ChR2^3^. Therefore, it is likely that our present study using chemogenetics was able to activate astrocytes under conditions more similar to physiological conditions than the previous study using optogenetics. In contrast, the activation of pons astrocytes strongly suppressed the onset of REM sleep, resulting in a significant decrease in the total time of REM sleep. The delta component of EEG during NREM sleep was significantly increased. Despite some differences, these results were generally in agreement with those of a previous study^22^. In addition, recent studies using chemogenetics demonstrated that different brain regions affect sleep/wakefulness states in different ways. The activation of astrocytes in the basal forebrain and lateral hypothalamus increases wakefulness and decreases NREM and REM sleep^24,28^. Furthermore, the activation of cortical astrocytes has been reported to increase sleep^21^. These results make it reasonable to assume that astrocytes contribute to sleep/wakefulness states differently depending on the brain region. Astrocytes have recently been demonstrated to be a heterogeneous population. Several types of astrocyte gene expression patterns have been identified through transcriptome analyses of various brain regions^29-32^. The heterogeneity of astrocytes may contribute to the different regulatory roles of astrocytes in the regulation of sleep/wakefulness.

### Possible mechanism of modulation of sleep/wakefulness states

What are the mechanisms by which activated astrocytes modulate sleep/wakefulness states and cortical oscillations? Regarding the pons, the brain region of hM3Dq expression in astrocytes coincides with the sublaterodorsal tegmental nucleus (SLD). Neurons located in the SLD are well-known to be REM-promoting neurons^33,34^. The activity of REM-promoting neurons in the SLD is strongly inhibited by gamma-amino butyric acid (GABA)ergic neurons in the dorsal part of the deep mesencephalic nucleus during wakefulness and NREM sleep^35,36^. It has been reported that activated astrocytes release glutamate, GABA, D-serine, adenosine triphosphate (ATP), and adenosine as gliotransmitters^37^. Astrocytes in the pons may release GABA, resulting in the suppression of REM-promoting neurons in the SLD. On the other hand, a previous study reported that inhibition of the increase in intracellular Ca^2+^ concentration in pons astrocytes decreases extracellular ATP concentration^22^, suggesting that pons astrocytes release ATP as gliotransmitters. The possibility that the ATP from astrocytes suppresses neurons in the SLD cannot be ruled out. Another possibility to consider is the involvement of microglia. Microglia have also been implicated in sleep/wakefulness regulation, and it is well known that they are activated during sleep deprivation^38-40^. We should consider the possibility that cytokines and reactive oxygen species released from astrocytes affect microglia, resulting in changes in sleep/wakefulness regulation.

The activation of astrocytes in the pons affected not only sleep/wakefulness states but also cortical oscillations. During NREM sleep, the delta component was significantly increased, whereas the theta, alpha, and beta components were significantly decreased. The homeostatic sleep drive, i.e., sleep need or sleep pressure, is represented by an increase in slow wave activity power^41,42^. Astrocytes regulate sleep pressure by releasing adenosine through soluble N-ethylmaleimide-sensitive factor attachment protein receptors^7,8^. Several reports showed that astrocyte Ca^2+^ signals express an accumulation of sleep pressure^9,43^. Thus, an increase in intracellular Ca^2+^ concentration in astrocytes may cause increased sleep pressure. The activation of hippocampal astrocytes, however, does not show a similar change in cortical oscillations. Additionally, a recent study proposed that activated astrocytes regulate sleep/wakefulness states independently of adenosine signals^22^. Taken together, there is presently insufficient lines of evidence to conclude whether increased astrocyte Ca^2+^ concentration leads to an increase in sleep pressure, and consequently, the delta component during NREM sleep. Further experiments are necessary to clarify this point.

The activation of hippocampal astrocytes significantly decreased the total time of wakefulness and increased the total time of sleep, but these were moderate alterations compared with those of the pons. It has been reported that the activity of hippocampal neurons changes according to sleep/wakefulness states^17,44,45^. However, there has been no lines of evidence to date that neurons are responsible for controlling sleep/wakefulness states. In addition, previous research has demonstrated that neurons in the brainstem are able to decode sleep and wakefulness states more adequately than those in the hippocampal region^13^. Considering these results, even though astrocytes in the hippocampus are activated and release gliotransmitters, they do not appear to have a substantial effect on sleep/wakefulness states. In addition, little effect on cortical oscillations was observed. Surprisingly, the theta component was not affected by the increase in astrocyte Ca^2+^ concentration in the hippocampus, despite it being demonstrated that astrocyte Ca^2+^ concentration in the cortex affects cortical oscillations^21,46^.

Our results as well as those of previous studies clearly indicate that astrocytes are involved in the regulation of sleep/wakefulness states. However, as Ca^2+^ concentrations in astrocytes were artificially increased in these studies, the extent to which they contribute to behavior under physiological conditions remains unclear. In addition, further studies are required to clarify the details regarding the type of gliotransmitters that are released from astrocytes, how these gliotransmitters regulate neural activity, and how they affect behavior.

## Acknowledgements

This work was supported in part by Precursory Research for Embryonic Science and Technology (PRESTO) (grant no.: JPMJPR1887) and Fusion Oriented Research for disruptive Science and Technology (FOREST) (grant no.: JPMJFR2047) from Japan Science and Technology Agency (JST), Shiseido Female Research Science Grant, and The Inamori Foundation Grant to T.T. Generous support from the FRIS CoRE, which is a shared research environment at Tohoku University, is acknowledged. We thank Professor Hiromu Tanimoto for helpful discussions, Sayuri Kobayashi for technical assistance, and Dr. Helena Akiko Popiel for English language editing of the manuscript.

